# Using photoaffinity labelling to study pantothenamide uptake in malaria parasites

**DOI:** 10.64898/2026.04.13.718166

**Authors:** Laura J. Akkerman, Solas Cassidy-Eulitz, Willem A. Velema, Taco W.A. Kooij

**Affiliations:** Department of Medical Microbiology, Radboudumc, Geert Grooteplein Zuid 28, Nijmegen, the Netherlands; Institute for Molecules and Materials, Radboud University, Heyendaalseweg 135, Nijmegen, the Netherlands

**Keywords:** malaria, *Plasmodium falciparum*, pantothenamide, photoaffinity labelling, drug uptake, gametocyte

## Abstract

Pantothenamides (PanAms) comprise a promising class of antimalarial compounds that kill asexual blood-stage *Plasmodium falciparum* parasites and block transmission. Intriguingly, the most advanced PanAm in drug development, MMV693183, is approximately 100 times more potent against female gametocytes than males. We hypothesized that this specificity is explained by a difference in PanAm uptake, which we studied using a PanAm-based photoaffinity labelling (PAL) probe. We successfully synthesized a probe that competed with MMV693183 in drug sensitivity assays, while the probe did not display high potency by itself. We observed no significant difference in median fluorophore-labelled probe signal intensity between male and female gametocytes, although there might be a difference in subcellular localization of the probe between the sexes. By combining PAL with affinity purification and mass spectrometry, we were not able to identify novel candidate PanAm transporters. We conclude that PAL provides evidence that differences in PanAm uptake do not underly differences in PanAm sensitivity between the gametocyte sexes.

## 1. Introduction

*Plasmodium* parasites cause more than 200 million cases of malaria every year (WHO, 2024). Drug resistance against front-line antimalarials is emerging and spreading (WHO, 2024), calling for the development of drugs with novel modes of action. Besides killing the asexual blood-stage parasites that cause malaria symptoms, these compounds should target the sexual blood-stage parasites or gametocytes that are responsible for malaria transmission to the mosquito (Burrows et al., 2013). One compound that meets these criteria is MMV693183 (Figure 1A) with IC_50_ ranges of 2.1-2.8 nM against asexual blood-stage *P. falciparum* parasites and 17.8-38.8 nM against gametocytes (de Vries et al., 2022). MMV693183 is a pantothenamide (PanAm), a synthetic analogue of pantothenate (vitamin B5, Figure 1B). Pantothenate is required for the biosynthesis of coenzyme A (CoA), which is involved in many processes including gene regulation, post-translational modifications, the tricarboxylic acid cycle, and fatty acid synthesis (de Vries et al., 2021). MMV693183 is metabolized in the CoA biosynthesis pathway to form CoA-MMV693183, an antimetabolite that inhibits the essential enzyme acetyl-CoA synthetase, killing the parasite (de Vries et al., 2022).

**Figure 1.**
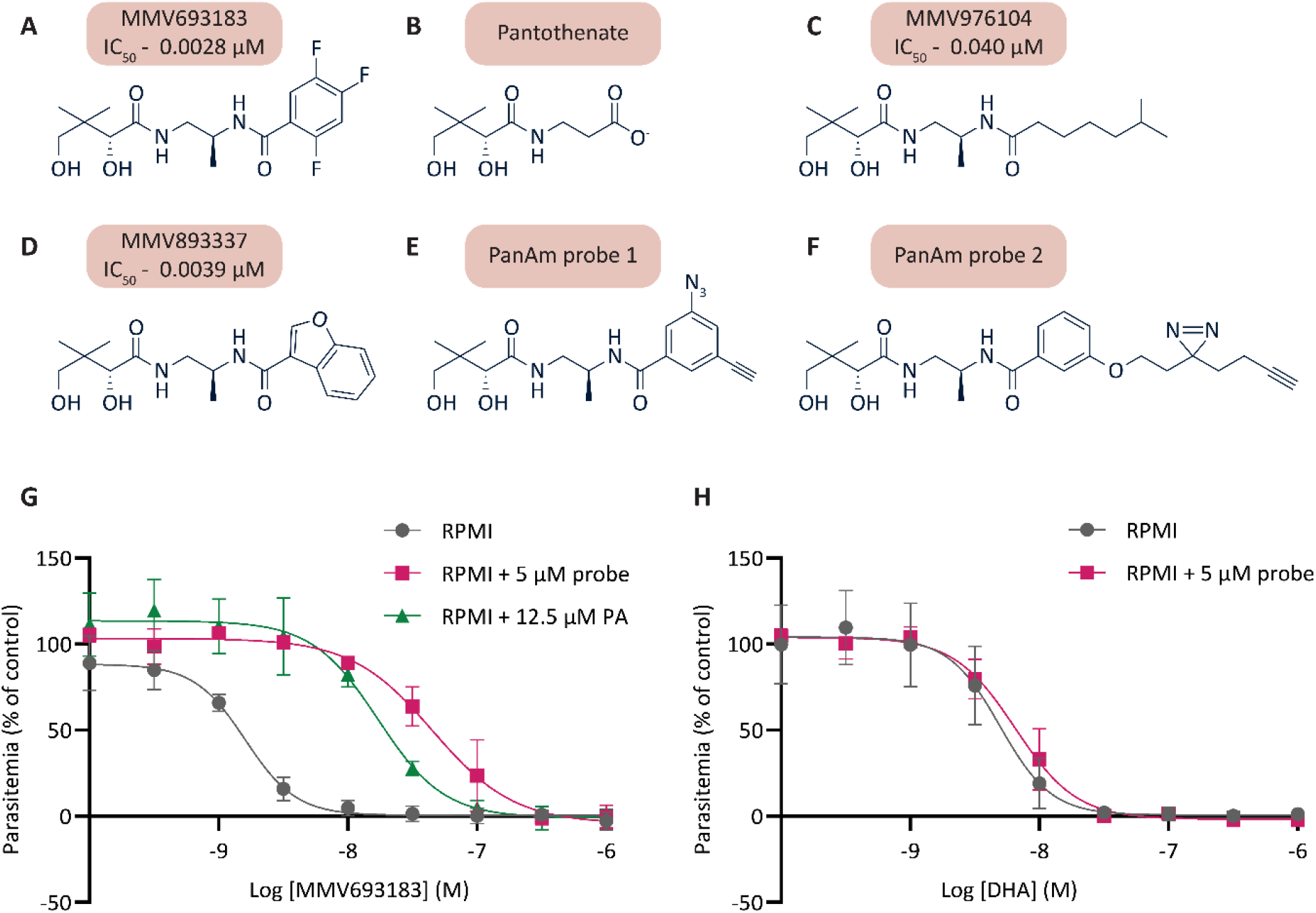
Development of a PanAm PAL probe. Chemical structures of A) MMV693183, B) pantothenate, C) MMV976104, D) MMV893337, E) PanAm probe 1, and F) PanAm probe 2. G/H) Dose-response curves of asexual blood-stage *P. falciparum* parasites for G) MMV693183 and H) dihydroartemisinin (DHA) in the presence of 5 µM PanAm probe 1, 12.5 µM pantothenic acid (PA), or no addition (RPMI only). Parasitaemia was normalized to DMSO-treated control for each condition. Average and standard deviation of three independent experiments are shown. The line corresponds to the best fit of a non-linear regression using a four-parameter model and the least squares method.

MMV693183 blocks human-to-mosquito transmission by killing gametocytes in human blood, rather than by affecting gametogenesis within the mosquito (de Vries et al., 2022). This means it requires a much shorter human plasma half-life than gametogenesis-inhibiting drugs, which need to be present exactly at the time of the mosquito bite, making it an attractive transmission-blocking compound (Birkholtz et al., 2022). Intriguingly, MMV693183 is approximately 100 times more potent against female gametocytes than male gametocytes (de Vries et al., 2022). We hypothesized that this difference in sensitivity is caused by a difference in uptake of PanAms between male and female gametocytes. It remains unknown how PanAms such as MMV693183 enter the malaria parasite but we expect that they follow the same route as pantothenate. While it has been established that there must be a proton-coupled pantothenate transporter on the parasite membrane (Saliba and Kirk, 2001), the identity of this protein remains elusive (Martin, 2020).

To study PanAm uptake in malaria parasites, we generated a PanAm probe to use for photoaffinity labelling (PAL). This technique is developed to identify molecular interactions between compounds and proteins (Singh et al., 1962; Mackinnon and Taunton, 2009; Smith and Collins, 2015) and has been successfully applied in many studies including in *P. falciparum* (Yahiya et al., 2023) and *Trypanosoma brucei* (Tulloch et al., 2017). The PAL probe harbours two functional groups: a UV-inducible reactive group to link the probe to any protein in its vicinity and a click-chemistry compatible group for labelling, for example with fluorophores or biotin. We combined PAL with microscopy to study uptake and localization of PanAms in gametocytes. We found no evidence that PanAm uptake is increased in female gametocytes, although the subcellular localization of the probe might be different between gametocyte sexes. By combining PAL with affinity purification and mass spectrometry, we aimed to identify new candidate PanAm transporters, which was unsuccessful. We conclude that PAL does not provide evidence for differential uptake of PanAms between gametocyte sexes, suggesting that there is another reason for PanAm specificity that remains unknown.

## 2. Materials & Methods

### Compounds

MMV693183 was synthesized previously (de Vries et al., 2022). Pantothenate was purchased as D-Pantothenic acid hemicalcium salt from Merck. See Supplemental Information for methods regarding synthesis, purification and analysis of PanAm probes.

### Parasite culture and replication assays

*P. falciparum* parasites were cultured in 5% human red blood cells (type O) (Sanquin), maintained in RPMI1640 medium supplemented with 25 mM HEPES, 25 mM NaHCO_3_, and 10% human type A serum (Sanquin) at 37 °C with 3% O_2_ and 4% CO_2_. Replication assays were performed according to a previously described protocol (Dery et al., 2015). In short, parasites were incubated with compounds for 72 hours in 394-well plates, followed by incubation with SYBR Green-containing lysis buffer for 1 hour, followed by measurement of fluorescence intensity on a BioTek Synergy 2 plate reader. Signals were normalized to a 0.1% DMSO control (100% growth) and a 1 µM dihydroartemisinin control (0% growth). Gametocytes were produced as reported previously (Graumans et al., 2024).

### Photoaffinity labelling

Cells (Figure 2: mixed asexual blood-stage (ABS) parasites, Figure 3: mature gametocytes) were washed once with RPMI1640 medium supplemented with 25 mM HEPES (incomplete medium), resuspended in incomplete medium with PanAm probe, transferred into a cell culture plate and incubated at 37 °C for 10 min. The plates (without lids) were then exposed to UV irradiation in a Stratagene UV Stratalinker 1800 for 3 times 5 min, shaking the plate to resuspend the cells in between. A pre-heated heat-block element was placed in the UV chamber to keep the samples at 37 °C throughout the irradiation step. Afterwards, the cells were transferred into microcentrifuge tubes, washed once with incomplete medium, and used for subsequent experiments.

**Figure 2.**
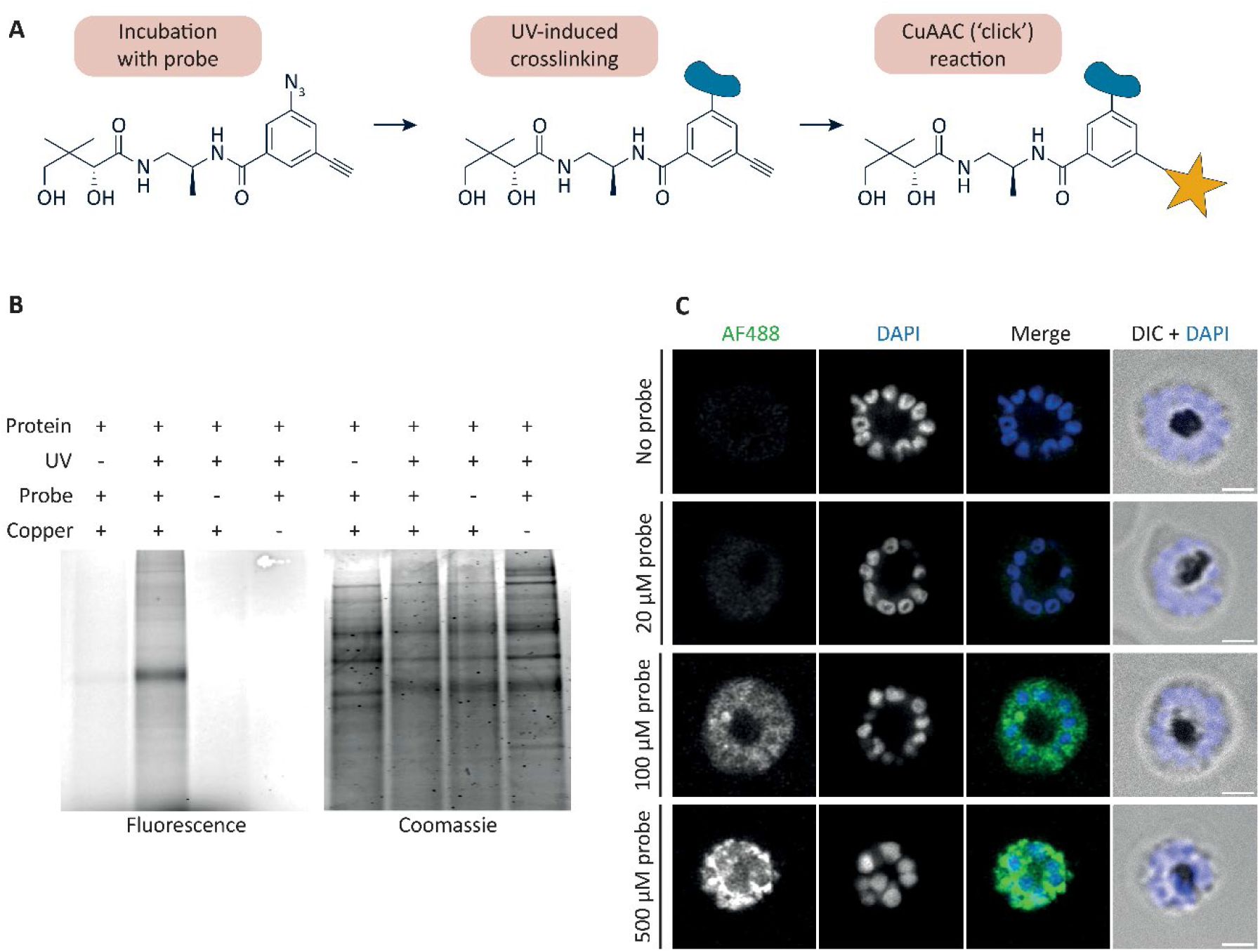
Validation of PAL and probe uptake. A) PAL workflow for SDS-PAGE or microscopy analysis. The blue shape represents any protein in the vicinity of the probe. The yellow star represents a fluorophore. CuAAC: copper(I)-catalysed azide-alkyne cycloaddition. B) SDS-PAGE analysis of parasite protein lysate after PAL procedure. C) Confocal PAL-microscopy images of asexual blood-stage *P. falciparum* parasites incubated with different probe concentrations. Probe is linked to Alexa Fluor 488 (AF488, green) and DNA is visualized with DAPI (blue). Scale bars: 2 µm. DIC: Differential Interference Contrast.

**Figure 3.**
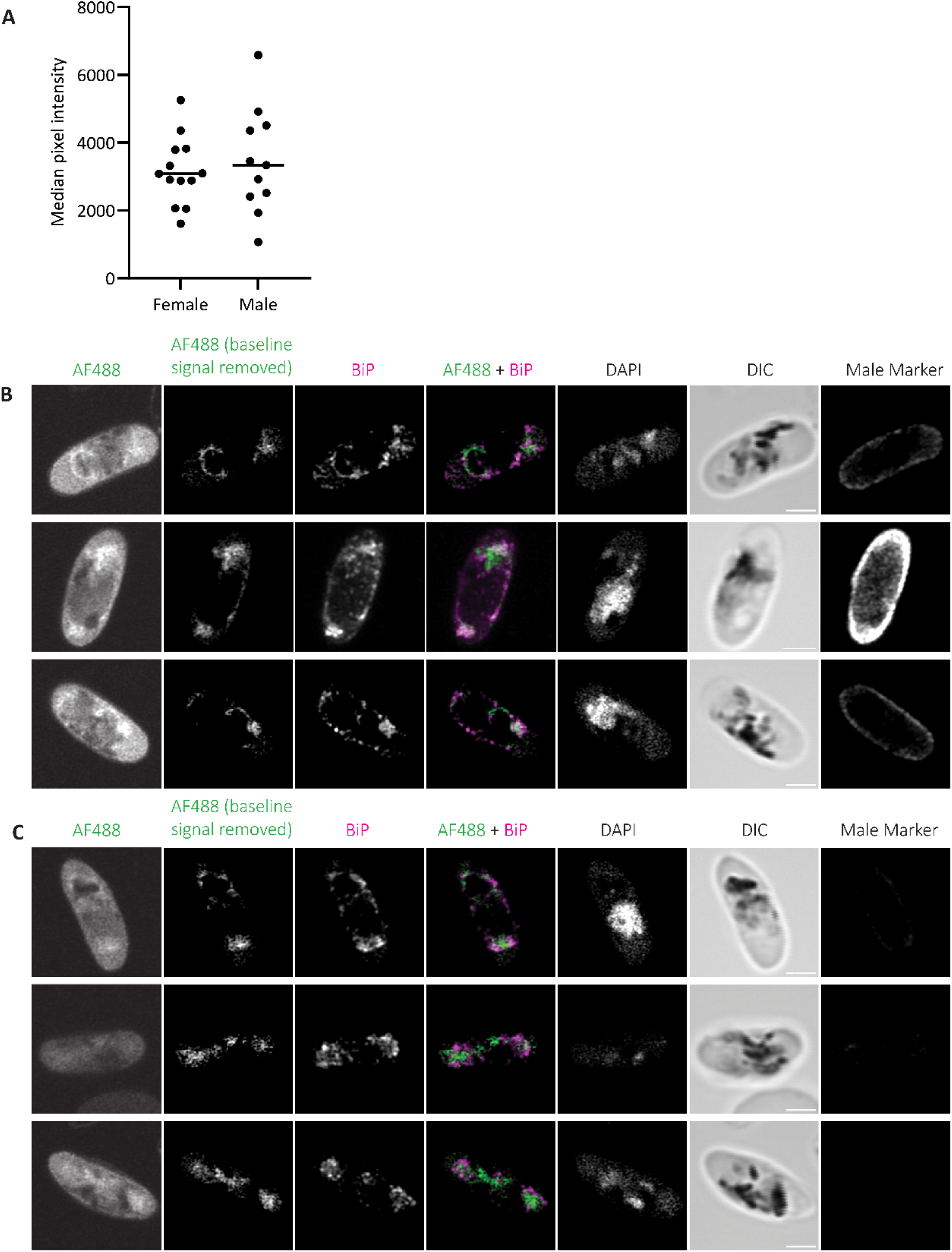
PanAm probe intensity and localization in gametocytes. A) Median probe signal intensity in female versus male gametocytes. B) Confocal immunofluorescence PAL-microscopy images of *P. falciparum* stage V gametocytes. Probe is linked to Alexa Fluor 488 (AF488, green), ER is stained with an anti-BiP antibody (magenta), and DNA is visualized with DAPI. The second image in each montage displays the probe signal after manual subtraction of the baseline signal. Scale bars: 2 µm. DIC: Differential Interference Contrast. C) Same as B for female gametocytes.

### (Immuno-)fluorescence staining and microscopy

After PAL, cells (Figure 2: mixed ABS parasites, Figure 3: mature gametocytes) were allowed to settle onto poly-L-lysine coated coverslips (Corning) for 20 min at 37 °C, followed by fixation for 20 min with 4% paraformaldehyde (Thermo Fisher Scientific) and 0.0075% glutaraldehyde (Thermo Fisher Scientific) in PBS (Tonkin et al., 2004). Cells were permeabilized with 0.1% Triton X-100 for 10 min and then incubated with 1% bovine serum albumin (BSA) (Sigma-Aldrich) in PBS for 5 min, washing with PBS in between the steps. Next, the coverslips were incubated with click reaction mixture: 0.5 mM copper(II) sulphate (Sigma-Aldrich), 5 mM sodium L-ascorbate (Sigma-Aldrich), 2.5 mM Tris(3-hydroxypropyltriazolylmethyl)amine (Sigma-Aldrich), and 40 µM AF 488 azide (Lumiprobe) in milli-Q for 30 min at room temperature (RT). Samples were then blocked with 3% BSA in PBS for 60 min, followed by incubation with primary antibody in 3% BSA in PBS for 60 min at RT, and then secondary antibody in PBS for 60 min at RT. Primary antibodies used were anti-SGDEDVDSDEL peptide (rabbit antiserum, BEI Resources) to detect the ER-resident chaperone BiP, and anti-LDH2 (rat antiserum, kindly provided by Dr Michael Delves) to detect male gametocytes (Sparkes et al., 2024). Secondary antibodies used were chicken anti-rabbit IgG (H+L) cross-adsorbed secondary antibody, Alexa Fluor™ 594 (Invitrogen) and goat anti-rat IgG (H+L) cross-adsorbed secondary antibody, Alexa Fluor™ 647 (Invitrogen). Coverslips were then incubated with 1 µM DAPI (Thermo Fisher Scientific) in PBS for 10 min, followed by mounting using Fluoromount-G mounting medium (Invitrogen). Imaging was performed on the Zeiss LSM900 Airyscan microscope with a 63X/1.4 oil objective. Airyscan processing of images was performed with Zeiss ZEN software, followed by further analysis (e.g. signal intensity quantification) using Fiji (Schindelin et al., 2012). For visualization, the same brightness/contrast settings were used for all images of the same channel within one figure, unless otherwise stated. For signal quantification, cell outlines were manually determined based on the DIC channel.

### Probe-protein complex affinity purification and mass spectrometry

After PAL on a mixed ABS parasite culture, RBCs were lysed by incubation with 0.06% saponin (Sigma-Aldrich) in PBS on ice for 10 min. Pellets were washed once with 0.06% saponin in PBS, twice with PBS, and then dissolved in 50 mM Tris-HCl solution (pH 8.8) with 1% SDS. Samples were homogenized by sonication, diluted to 0.5% SDS, centrifuged at >16,000 g for 20 min at 4°C. The supernatant was transferred to a new tube and the protein concentration was determined by Pierce BCA Protein Assay Kit (Thermo Fisher Scientific) according to the manufacturers instruction . For each sample, 120 µg protein was incubated with click reaction mixture: 0.5 mM copper(II) sulphate (Sigma-Aldrich), 5 mM sodium L-ascorbate (Sigma-Aldrich), 2.5 mM Tris(3-hydroxypropyltriazolylmethyl)amine (Sigma-Aldrich), and 100 µM azide-PEG3-biotin conjugate (Sigma-Aldrich) in milli-Q for 30 min at RT. Zeba spin desalting columns (7K MWCO) (Thermo Fisher Scientific) were used according to manufacturer’s instructions to remove redundant biotin azide and other small-molecule contaminants. Using MagReSyn streptavidin mass spectrometry beads, biotin-probe-protein complexes were purified followed by on-bead reduction, alkylation, and digestion according to manufacturer’s instructions.

Tryptic digests were analyzed by nanoflow liquid chromatography (Evosep One, Evosep Biosystems) coupled online to a trapped ion mobility spectrometry – quadrupole time-of-flight mass spectrometer (timsTOF Pro2, Bruker Daltonics) *via* a nanoflow electrospray ionization source (CaptiveSprayer, Bruker Daltonics). Tryptic peptides were separated by C18 reversed phase liquid chromatography (Evosep EV1137 30SPD performance column; 150 mm length x 0.150 mm internal diameter, 1.5 µm C18AQ particles) using the pre-programmed 30 samples per day (30SPD) Evosep One method. The mass spectrometer was operated in positive ionization mode using the default data dependent acquisition – Parallel Accumulation SErial Fragmentation (dda-PASEF) instrument method: mass range: 100-1700 m/z, mobility range: 0.6-1.6 1/K_0_, accumulation time: 100 ms, ramp time: 100 ms, PASEF cycles: 10, dynamic exclusion 0.4 min). Acquired spectra were streamed directly to PaSER box (v2024, Bruker Daltonics) for protein identification and label-free quantitation against the combined Uniprot reviewed human and Uniprot *Plasmodium falciparum* (isolate 3D7) protein sequences (downloaded Jan 2024) using the following settings: ProLuCID search engine, 20 ppm precursor mass tolerance, 30 ppm fragment ion mass tolerance, strict tryptic cleavage, maximum of 2 missed cleavages, carbamidomethyl (C) as fixed modification, deamidation (NQ), and oxidation (M) as variable modifications. Individual search results were combined and validated using DTA select (Tabb et al., 2002) with TIMScore enabled to achieve ≤1% false discovery rate at protein level. Each sample was prepared and measured in duplicate. Hits that were only detected in one of the two duplicates for the probe-treated samples were excluded.

3. Results

### 3.1 Synthesis of a PanAm PAL probe

The most advanced PanAm in development as an antimalarial drug is MMV693183 (de Vries et al., 2022) (Figure 1A), a derivative of pantothenate (Figure 1B). To design a PanAm probe for PAL, we focused on making modifications around the fluorinated aromatic ring in MMV693183, since these alterations are tolerated whilst retaining potency. This is demonstrated by MMV976104 and MMV893337, which both display nanomolar activity against asexual blood-stage parasites despite their distinct chemical structure (Figure 1C-D) (Schalkwijk et al., 2019). We successfully synthesized two probes, both containing an alkyne handle for copper(I)-catalysed azide-alkyne cycloaddition (CuAAC) reactions and a photoactivatable group, which was a phenyl azide for probe 1 (Figure 1E) and a diazirine for probe 2 (Figure 1F).

We measured activity of the probes against asexual blood-stage *P. falciparum* parasites *in vitro*. Against our expectations, both probes were found to be much less potent than MMV693183, with IC_50_ values above 1 µM (Supplementary Figure 1A). For all subsequent experiments, we used probe 1, whose chemical structure closer resembles that of more active pantothenamides. We hypothesized that the probe might not be completely metabolized by the CoA biosynthesis pathway into its active form, explaining its low potency. Therefore, we performed a competition drug sensitivity assay, in which the parasite’s sensitivity to MMV693183 was determined in the presence of a fixed amount of probe or pantothenic acid, which is a known PanAm competitor and served as a positive control (Schalkwijk et al., 2019). Parasites displayed a lower sensitivity to MMV693183 in the presence of 5 µM probe, similar to the reduced sensitivity in the presence of 12.5 µM pantothenic acid (Figure 1G). As a negative control, we tested whether the addition of probe affected the parasite’s sensitivity to dihydroartemisinin, which was not the case (Figure 1H). PanAm-probe competition was concentration-dependent: the more probe was added, the less sensitive to MMV693183 the parasites became (Supplementary Figure 1B). We concluded that the probe competes with MMV693183 for uptake and/or metabolic processing and is suitable to study PanAm uptake.

### 3.2 Validation of probe-protein crosslinking and uptake in live parasites

First, we tested if the PanAm probe could be crosslinked to proteins in asexual blood-stage malaria parasites *in vitro* (Figure 2A). We incubated infected red blood cells (iRBCs) with the probe, followed by exposure to UV radiation to induce formation of a reactive nitrene intermediate, which immediately and covalently binds any protein in its surroundings. We then collected the cells and lysed them, after which we linked the protein-probe complexes to a fluorophore-azide using a CuAAC reaction. Analysis of the lysate by SDS-PAGE confirmed that fluorophore-probe-protein complexes were present (Figure 2B). Spontaneous binding of probe in the absence of UV, as well as spontaneous binding of fluorophore-azide in the absence of probe or the catalyst Cu(I), was negligible.

Next, we combined the photo-affinity labelling procedure with microscopy. To do this, we incubated iRBCs with probe and exposed to UV radiation, after which they were settled on coverslips, fixed, and permeabilized. Afterwards, we ‘clicked on’ the Alexa Fluor 488 azide, followed by antibody and dye staining. We successfully visualized the probe and observed that increasing probe concentration during the incubation step resulted in increased signal intensity (Figure 2C). These initial experiments confirmed that photo-affinity labelling with the PanAm probe can be used to study PanAm uptake in live malaria parasites.

### 3.3 Studying uptake of pantothenamides by gametocytes

Male gametocytes are approximately 100-fold less sensitive to MMV693183 than female gametocytes (de Vries et al., 2022). To test whether the insensitivity of male gametocytes to PanAms can be attributed to a reduced uptake, we studied uptake of our probe in gametocytes with microscopy. To this end, we incubated stage V gametocytes with 100 µM probe and performed PAL, followed by immunofluorescence staining with a male-specific antibody to determine gametocyte sex. Although there was variability in the probe-derived signal between gametocytes, there was no significant difference between median probe signal intensities in male versus female gametocytes (Figure 3A). Upon closer inspection, the probe signal seemed to be more intense within certain intracellular structures of the gametocytes. In male gametocytes, the brightest probe signal appeared to localize to the endoplasmic reticulum (ER), which was stained with an antibody against the abundant ER-resident chaperone BiP, a member of the heat-shock protein 70 family (Figure 3B). We did not observe this colocalization in female gametocytes (Figure 3C).

### 3.4 Mass spectrometry-based identification of probe-protein complexes

Next, we aimed to identify PanAm transporter candidates by combining PAL with mass spectrometry-based proteomics. To this end, we performed PAL as before but ‘clicked’ a biotin azide molecule to the probe-protein complexes to enable affinity purification followed by protein digestion and mass spectrometry (Figure 4A). In a preliminary experiment, we treated asexual blood-stage parasites with 0, 1 or 10 µM probe for 10 minutes before crosslinking, expecting that specific interactions between the probe and its transporter or downstream interactors would reflect in a concentration-dependent enrichment. From the 70 proteins that were more than two times enriched in either sample, only 7 hits overlapped (Figure 4B-C). None of these had 2 or more predicted transmembrane domains, which would be expected in a transporter. The proteins of the CoA biosynthesis pathway, which we expected to metabolize the probe, were not detected in any of the samples. One potentially interesting hit was multidrug resistance protein 2, which was enriched in the 10 µM sample but not in the 1 µM sample. However, since this protein is dispensable and associated with heavy metal transport (Rosenberg et al., 2006; Rosental et al., 2012; van der Velden et al., 2015), we deemed it an unlikely candidate for PanAm transport. Overall, we concluded that our approach was not suitable to identify interaction partners of the probe.

**Figure 4.**
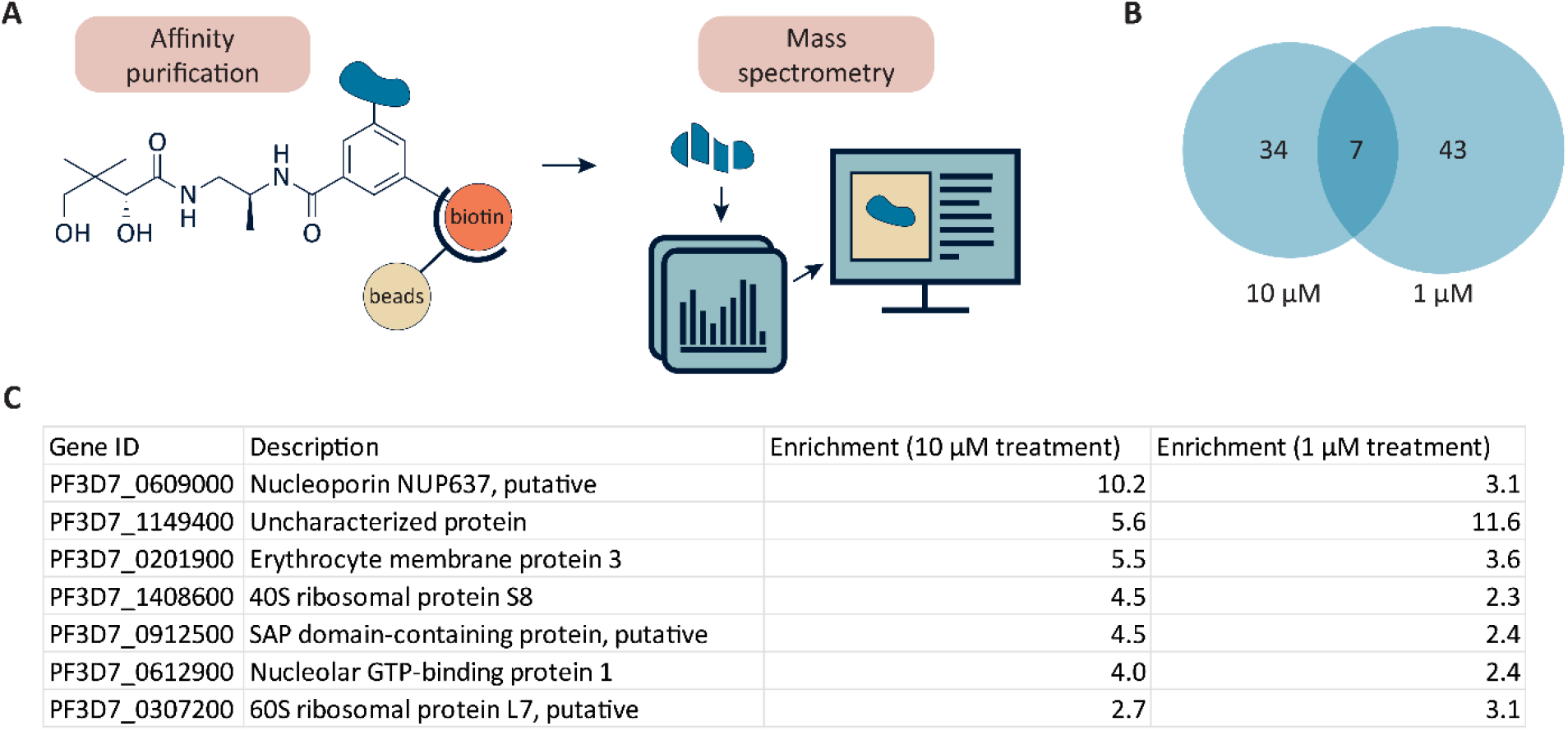
Identification of probe interaction partners. A) Workflow of PAL combined with affinity purification and mass spectrometry. B) Venn diagram of hits that were >2 times enriched when parasites were incubated with 1 or 10 µM probe compared to no probe in a preliminary experiment. C) Hits that were >2 times enriched upon treatment with 1 and 10 µM probe compared to no probe in a preliminary experiment.

## 4. Discussion

The development of transmission-blocking compounds such as PanAms into novel drugs is critical to stop the spread of malaria (Burrows et al., 2013). We aimed to characterize uptake of PanAms using PAL combined with microscopy and affinity purification followed by mass spectrometry. We successfully synthesized a PanAm probe and confirmed that it is taken up by asexual blood-stage parasites and gametocytes, and that it competes with MMV693183, indicating that it can be used as a tool to study PanAm uptake. Based on the observed 100-fold higher sensitivity of female over male gametocytes, we hypothesized that female gametocytes import PanAms more efficiently than males. However, we observed no significant difference in median probe signal intensities between male and female gametocytes. Nevertheless, we demonstrate that PAL combined with microscopy is a powerful technique to study uptake and subcellular localization of small molecules. The photoactivatable group and the click chemistry-compatible group are relatively small, making this approach more interesting than directly attaching a fluorophore to the molecule of interest. Furthermore, co-staining of the samples with antibodies allows for colocalization of the probe with a structure of interest. In the future, this approach could be used to study subcellular localization of more drugs, vitamins, and other molecules of interest.

It remains unclear why PanAms specifically kill female and not male gametocytes. Rather than a difference in drug uptake, another distinction between male and female gametocytes probably explains PanAm specificity. For example, the PanAm-affected pathway(s) might be more critical for females than males, but these downstream effects are not entirely understood (de Vries et al., 2021). One potential PanAm-affected pathway is histone acetylation, which is known to be crucial for gene regulation in gametocytes (Ngwa et al., 2017). However, other drugs that affect histone modifications are one order of magnitude more potent against male gametogenesis than female gametogenesis (Malmquist et al., 2015; Matthews et al., 2020). Alternatively, there might be differences in (pro)drug metabolism between male and female gametocytes, but PanAm metabolism is only studied in asexual blood-stage parasites so far (de Vries et al., 2022). It could also be that the subcellular distribution of the active PanAm metabolite is different between male and female gametocytes, resulting in lower levels of inhibitor at the target site. In line with this hypothesis, we observed a possible difference in localization of our PanAm probe between gametocyte sexes, but this observation should be replicated with a more potent probe to draw a strong conclusion.

PAL followed by mass spectrometry-based identification of probe-protein complexes did not yield any new candidate PanAm transporters. While PAL has been used to identify transporters in the past (e.g. (Kurosawa et al., 2022)), the chance of success depends on the specificity of the probe-protein interactions, sensitivity to detect these interactions, and the timing of the experiment. Specificity to detect specific probe-protein interactions over non-specific interactions (i.e. random proximity to other proteins) depends on the strength/duration of the interaction, and probe-transporter binding might be too transient to detect. Additionally, the proteins that interact with the probe might be lowly expressed, challenging the detection limits of the approach. It is also possible that most of the probe had already been taken up within the 10-minute incubation period after which we started crosslinking. However, if timing (rather than specificity and sensitivity) was the only problem, we would still expect to detect downstream probe-interacting proteins, such as the proteins of the CoA biosynthesis pathway, which we did not. In future studies, the PanAm transporter should be identified using an alternative method. Assuming that PanAms and pantothenate enter through the same transporter, one could perform a targeted study of putative pantothenate transporters. Unfortunately, these are limited: a thorough review (Martin, 2020) yielded 2 main candidates: MCP1 and UMF, which are both hypothesized to localize to the apicoplast rather than the parasite membrane based on studies in *Toxoplasma* (Chen et al., 2024; Dong et al., 2024) and, for UMF, biotin proximity labelling in *P. falciparum* (Boucher et al., 2018). To identify new transporter candidates, one could perform an unbiased *in vitro* search for putative pantothenate/PanAm transporters, for example using thermal proteome profiling (Reinhard et al., 2015).

The probes generated in this study displayed limited potency against asexual blood-stage parasites. Given that PanAms are prodrugs that need to be activated by enzymes of the CoA biosynthesis pathway (de Vries et al., 2021), it could be that the probe is not efficiently processed into its active form, that its active form is processed further, or that its active form binds its target less well. A more potent probe would be a useful tool to study the downstream effects of PanAms, although its design is complicated by the fact that the structure-activity relationship of PanAms is not entirely understood (de Vries et al., 2022). The subcellular localization of a potent PanAm probe could indicate the most likely mode of action: if potent PanAms accumulate, for example, in the nucleus, the most likely mode of action would be to disrupt gene regulation. Furthermore, studying the interaction partners of a potent PanAm probe could shed more light on the mechanism of action of PanAms. Specifically, while the inhibition of acetyl-CoA synthetase by PanAm metabolites has been confirmed, other enzymes such as the synthetic lethal pair of alpha-ketoglutarate dehydrogenase and mitochondrial pyruvate dehydrogenase also produce acetyl-CoA (Nair et al., 2023) and might therefore be inhibited as well. Taken together, more potent PanAm probes could provide more answers on mode and mechanism of action of PanAms.

## Author contributions

**L.J. Akkerman:** Conceptualization, Formal analysis, Funding acquisition, Investigation, Methodology, Supervision, Visualization, Writing – original draft, Writing – review & editing. **S. Cassidy-Eulitz:** Investigation, Methodology, Writing – review & editing. **T.W.A. Kooij:** Conceptualization, Funding acquisition, Project administration, Supervision, Writing – review & editing. **W.A. Velema:** Conceptualization, Funding acquisition, Supervision, Writing – review & editing.

## Acknowledgements

We thank Nicholas Proellochs for providing stage V gametocyte samples. This work was supported by the Radboud University Medical Center (Radboudumc Individual PhD for Masters round 2021 to LJA). Proteomics measurements were performed by the Radboud Technology Center for Mass Spectrometry supported by the Netherlands X-omics Initiative partially funded by NWO (project 184.034.019).

