## Supplemental Information for "Using photoaffinity labelling to study pantothenamide uptake in malaria parasites"

### Supplemental Figures

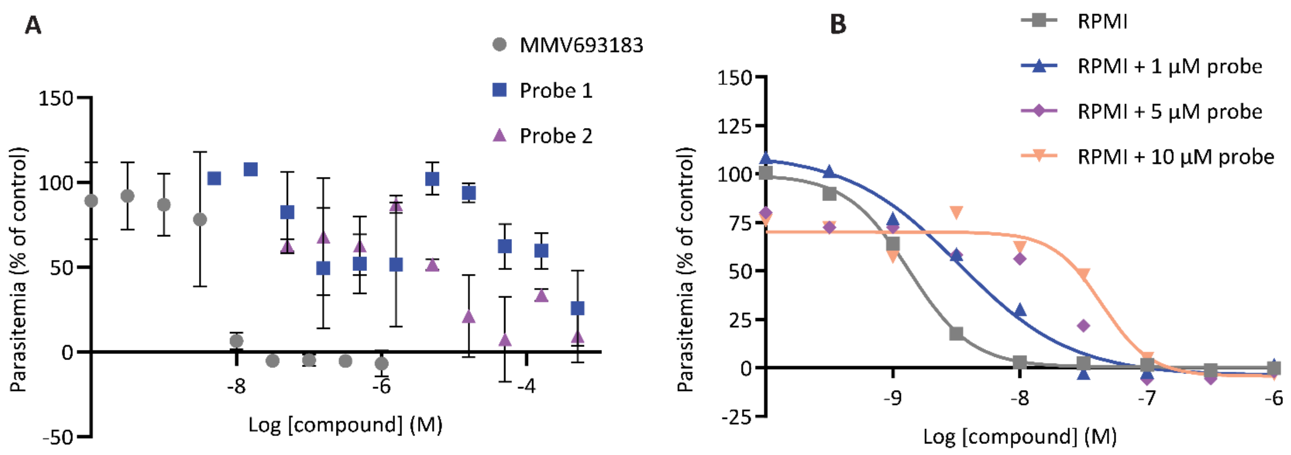


***Figure S1 – Development of a PanAm PAL probe.*** *A) Dose-response measurements of asexual P. falciparum parasites for PAL probe 1 and 2. Average values and SEM ranges are based on two independent experiments. B) Dose-response curves of asexual* P. falciparum *parasites for MMV693183 with different fixed concentration of probe 1 added to the medium.*

### Synthesis, purification and analysis of PanAm probes

Commercially available chemicals were purchased and used as received without further purification. Analytical thin-layer chromatography (TLC) was performed using glass plates pre-coated with silica gel (0.20 mm, 60 Å pore size) and fluorescent indicator UV254 (Merck) and TLC-aluminium sheets pre-coated with silica gel (0.20 mm, 60 Å pore size) and fluorescent indicator UV254 (Merck). TLC plates were visualized by exposure to ultraviolet light (UV) and/or using potassium permanganate (KMnO4) solution followed by brief heating with a heat gun (10–15 s). Normal phase flash-column chromatography was carried out using Biotage® Isolera™ Systems. NMR spectra were recorded at ~23 °C on a Bruker Avance III 500 MHz. The 1H and the 13C NMR chemical shifts were given in parts per million (ppm) relative to the residual signals of the deuterated solvents. Carbon types were determined from APT and CPD 13C NMR. Mass spectra were recorded on a Single-Quad Thermo ISQ with UltiMate 3000 autosampler and LC-MS (ESI) 95% PDA (254nm) spectrometer. Instrumental method: 10 min run + MS (0 – 95 MilliQ-ACN) Accucore™ C18 10min.

#### Synthesis of probe 1

2,5-dioxopyrrolidin-1-yl 3-azido-5-ethynylbenzoate – Intermediate 2

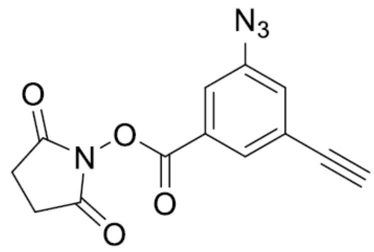

A solution of 3-azido-5-ethynylbenzoic acid (75 mg, 1 Eq, 0.40 mmol) **(Intermediate 1**, synthesized as previously described by Pošta, Soós & Beier, *Tetrahedon*, 2016), 1-hydroxypyrrolidine-2,5-dione (69 mg, 1.5 Eq, 0.60 mmol) and EDC (115 mg, 1.5 Eq, 600 µmol) in DCM (4.9 mL) were stirred at room temperature overnight and analysed via TLC. The solvent was evaporated *in vacuo*, diluted with EtOAc, and washed with water, $NaHCO_3$, brine and dried over $MgSO_4$. The organics were combined and concentrated to obtain the crude product 2,5-dioxopyrrolidin-1-yl 3-azido-5-ethynylbenzoate (118 mg, 415 µmol, quantitative) as a yellow/brown solid.

**Rf:** 0.41 (1:1 EtOAc in Heptane) **1H NMR:** (500 MHz, CHLOROFORM-D) δ 7.99 (t, J = 1.5 Hz, 1H), 7.72 (dd, J = 2.3, 1.5 Hz, 1H), 7.39 (dd, J = 2.3, 1.4 Hz, 1H), 3.19 (s, 1H), 2.93 – 2.87 (m, 4H). **13C NMR** (126 MHz, CHLOROFORM-D) δ 168.95, 160.59, 141.66, 130.32, 128.20, 127.26, 125.00, 120.98, 80.99, 80.20, 25.75.

tert-butyl (*S*)-(2-(3-azido-5-ethynylbenzamido)propyl)carbamate – Intermediate 3

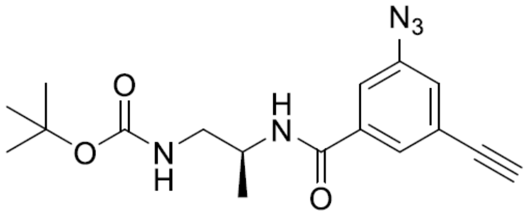

A solution of 2,5-dioxopyrrolidin-1-yl 3-azido-5-ethynylbenzoate (120 mg, 1 Eq, 422 µmol) **(Intermediate 2)**, tert-butyl *(S)*-(2-aminopropyl)carbamate (80.9 mg, 1.1 Eq, 464 µmol) and $Et_3N$ (128 mg, 177 µL, 3 Eq, 1.27 mmol) were dissolved in DCM. The reaction mixture was stirred at room temperature overnight. A TLC and crude NMR confirmed the reaction had got to completion. The solvent was removed under vacuum and the crude product was dissolved in EtOAc and poured into water. The aqueous fraction was washed with EtOAc, and the combined organic fractions were washed with a saturated aqueous solution of $NaHCO_3$, brine, dried over $MgSO_4$ and concentrated in vacuo. Purification using flash silica chromatography (20% → 45% EtOAc in heptane) afforded pure product tert-butyl (*S*)-(2-(3-azido-5-

ethynylbenzamido)propyl)carbamate (182 mg, 530 µmol, quantitative) as a white solid. Solvent impurities present.

**Rf:** 0.70 (1:1 EtOAc in Heptane) **1H NMR** (500 MHz, CHLOROFORM-D) δ 7.67 (t, J = 1.5 Hz, 1H), 7.53 (t, J = 1.9 Hz, 1H), 7.44 (d, J = 6.6 Hz, 1H), 7.19 (dd, J = 2.2, 1.3 Hz, 1H), 5.00 (t, J = 6.5 Hz, 1H), 4.15 (q, J = 3.1 Hz, 1H), 3.33 (ddd, J = 14.6, 8.8, 7.5 Hz, 1H), 3.27 (s, 1H), 3.22 (ddd, J = 14.8, 5.6, 3.6 Hz, 1H), 3.10 (s, 1H), 1.41 (s, 9H).

(*S*)-*N*-(1-aminopropan-2-yl)-3-azido-5-ethynylbenzamide – Intermediate 4

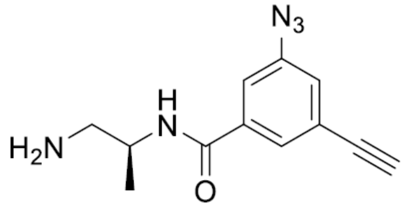

Tert-butyl (*S)*-(2-(3-azido-5-ethynylbenzamido)propyl)carbamate (180 mg, 1 Eq, 524 µmol) **(Intermediate 3)** was dissolved in 5mL of 4N HCL in Dioxane (excess). The reaction mixture was stirred at room temperature and checked via TLC. Once complete the reaction mixture was concentrated in vacuo to obtain crude product *(S)*-*N*-(1-aminopropan-2-yl)-3-azido-5-ethynylbenzamide (135 mg, 555 µmol, quantitative) as white solid.

**Rf:** 0.04 (5:1 EtOAc in Heptane) **1H NMR** (500 MHz, METHANOL-D4) δ 7.75 (t, J = 1.5 Hz, 1H), 7.56 (t, J = 1.9 Hz, 1H), 7.22 (dd, J = 2.3, 1.3 Hz, 1H), 4.40 – 4.29 (m, 1H), 3.43 (s, 1H), 3.07 (d, J = 6.5 Hz, 2H), 1.34 – 1.27 (m, 3H).

3-azido-*N*-((*S*)-1-((*R*)-2,4-dihydroxy-3,3-dimethylbutanamido)propan-2-yl)-5- ethynylbenzamide – Probe 1

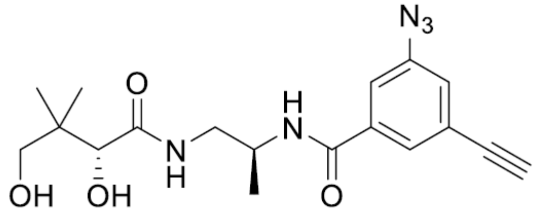

A solution of (*S*)-*N*-(1-aminopropan-2-yl)-3-azido-5- ethynylbenzamide (135 mg, 1 Eq, 555 µmol) **(Intermediate 4)**, (*S*)-3-hydroxy-4,4-dimethyldihydrofuran-2(3H)-one (79.4 mg, 1.1 Eq, 610 µmol) and $Et_3N$ (168 mg, 232 µL, 3 Eq, 1.66 mmol) in EtOH (9 mL) were added to a microwave flask and stirred over the weekend at 80 °C. The reaction was monitored using TLC. The reaction mixture was concentrated in vacuo and purified using flash silica chromatography (0% → 10% MeOH in DCM) to afforded pure product 3-azido-*N*-((*S*)-1-((*R*)-2,4-dihydroxy-3,3-dimethylbutanamido)propan-2-yl)-5-ethynylbenzamide (68 mg, 0.18 mmol, 33 %) as an off white solid.

**Rf:** 0.73 (1:10 MeOH in DCM) **1H NMR** (500 MHz, METHANOL-D4) δ 7.69 (t, J = 1.5 Hz, 1H), 7.50 (t, J = 1.9 Hz, 1H), 7.24 (dd, J = 2.2, 1.4 Hz, 1H), 4.29 – 4.18 (m, 1H), 3.89 (s, 1H), 3.64 (s, 1H), 3.44 – 3.28 (m, 5H), 1.22 (d, J = 6.7 Hz, 3H), 0.82 (d, J = 4.5 Hz, 6H). **13C NMR** (126 MHz, METHANOL-D4) δ 175.43, 166.49, 141.15, 136.52, 127.01, 124.54, 124.37, 118.15, 81.34, 79.41, 76.06, 69.16, 53.48, 46.57, 43.59, 29.40, 17.06. **MS** (ESI) (m/z) calculated for $[M+H]^+$:374.18, found 373.83; calculated for $[M+Na]^+$: 396.16, found 396.14.

#### Synthesis of probe 2

methyl 3-(2-(3-(but-3-yn-1-yl)-3H-diazirin-3-yl)ethoxy)benzoate – Intermediate 5

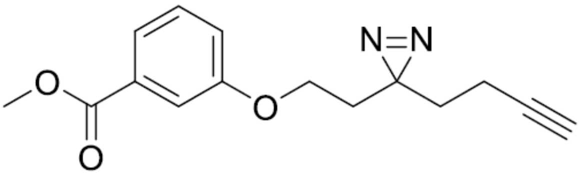

3-(but-3-yn-1-yl)-3-(2-iodoethyl)-3H-diazirine (10 mg, 10 µL, 1 Eq, 40 µmol) was added to a stirred solution of methyl 3-hydroxybenzoate (4.6 mg, 0.75 Eq, 30 µmol) and potassium

carbonate (11 mg, 2 Eq, 81 µmol) in Acetone (1.5 mL). The reaction mixture was refluxed overnight and analysed and TLC. The crude product was extracted with EtOAc (3 x 10 mL), washed with water, washed with brine, dried over MgSO4, and concentrated *in vacuo*. Purification via silica flash-column chromatography (0% → 40% EtOAc in heptane) to yield pure product methyl 3-(2-(3-(but-3-yn-1-yl)-3H-diazirin-3-yl)ethoxy)benzoate (4 mg, 0.01 mmol, 40 %) as a white solid.
**Rf:** 0.78 **(**5:2 EtOAc in Heptane) **1H NMR** (500 MHz, CHLOROFORM-D) δ 7.63 (dt, J = 7.7, 1.3 Hz, 1H), 7.52 (dt, J = 2.8, 1.4 Hz, 1H), 7.33 (t, J = 7.9 Hz, 1H), 7.08 (ddd, J = 8.2, 2.7, 1.0 Hz, 1H), 3.90 (s, 3H), 3.86 (s, 1H), 2.05 (td, J = 7.5, 2.6 Hz, 2H), 1.98 (t, J = 2.7 Hz, 1H), 1.89 (t, J = 6.2 Hz, 2H), 1.73 (t, J = 7.5 Hz, 2H).

<u>3-(2-(3-(but-3-yn-1-yl)-3H-diazirin-3-yl)ethoxy)benzoic acid – Intermediate 6</u>

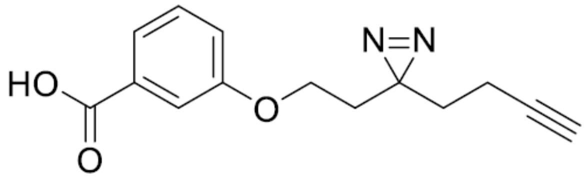

Potassium Hydroxide (52 mg, 10 Eq, 0.92 mmol) was dissolved in water (1.5 mL) and added to a stirred solution of methyl 3-(2-(3-(but-3-yn-1-yl)-3H-diazirin-3-yl)ethoxy)benzoate (25 mg, 1 Eq, 92 µmol) **(Intermediate 5)** in THF (4.5 mL) and refluxed overnight at 70 °C. The reaction mixture was slowly quenched with HCl and washed with DCM (x3). The combined organic layers were dried over $MgSO_4$, filtered, and concentrated *in vacuo*. The crude was purified via silica flash-column chromatography (0% → 40% EtOAc in heptane) to yield pure product 3-(2-(3-(but-3-yn-1-yl)-3H-diazirin-3-yl)ethoxy)benzoic acid (20 mg, 77 µmol, 84 %) as a white solid.
**Rf:** 0.30 **(**1:1 EtOAc in Heptane) **1H NMR** (500 MHz, CHLOROFORM-D) δ 7.71 (dt, J = 7.8, 1.3 Hz, 1H), 7.58 (dd, J = 2.7, 1.5 Hz, 1H), 7.37 (t, J = 7.9 Hz, 1H), 7.14 (ddd, J = 8.2, 2.7, 1.0 Hz, 1H), 3.88 (t, J = 6.2 Hz, 2H), 2.07 (td, J = 7.5, 2.7 Hz, 2H), 1.99 (t, J = 2.6 Hz, 1H), 1.91 (t, J = 6.2 Hz, 2H), 1.74 (t, J = 7.5 Hz, 2H).

<u>tert-butyl ((*S*)-1-((*R*)-2,4-dihydroxy-3,3-dimethylbutanamido)propan-2-yl)carbamate – Intermediate 7</u>

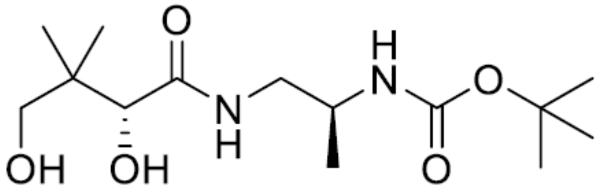

To a microwave flask was added tert-butyl (*S*)-(1- aminopropan-2-yl)carbamate (250 mg, 1 Eq, 1.43 mmol), *(R)-* 3-hydroxy-4,4-dimethyldihydrofuran-2(3H)-one (205 mg, 1.1 Eq, 1.58 mmol) and $Et_3N$ (218 mg, 300 µL, 1.5 Eq, 2.15 mmol) in EtOH (4 mL) and stirred over the weekend at 80 °C. The reaction was monitored via TLC and stained with $KMnO_4$. The crude (yellow oil) was purified via silica flash-column chromatography (0 → 10% MeOH in DCM) 25 g silica column, eluting with 7% MeOH to yield purified product tert-butyl ((*S*)-1- ((*R*)-2,4-dihydroxy-3,3-dimethylbutanamido)propan-2-yl)carbamate (281 mg, 923 µmol, 64.3 %).
**Rf:** 0.58 **(**1:10 MeOH in DCM) **1H NMR** (500 MHz, CHLOROFORM-D) δ 7.32 (t, J = 6.1 Hz, 1H), 5.05 (d, J = 7.8 Hz, 1H), 4.70 – 4.65 (m, 1H), 4.05 (s, 1H), 3.98 (d, J = 4.9 Hz, 1H), 3.71 (p, J = 6.9 Hz, 1H), 3.43 (d, J = 4.8 Hz, 2H), 3.32 – 3.16 (m, 3H), 1.37 (s, 9H), 1.10 (d, J = 6.7 Hz, 3H), 0.94 (s, 3H), 0.87 (s, 3H).

(*R*)-*N*-((*S*)-2-aminopropyl)-2,4-dihydroxy-3,3-dimethylbutanamide – Intermediate 8

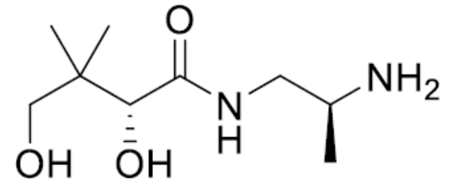

tert-butyl ((*S*)-1-((*R*)-2,4-dihydroxy-3,3-dimethylbutanamido)propan-2-yl)carbamate (281 mg, 1 Eq, 923 µmol) **(Intermediate 7)** was dissolved in 5 mL of 4N HCL in Dioxane (excess). The reaction mixture was stirred at room temperature and checked via TLC. Once complete the reaction mixture was concentrated *in vacuo* to obtain crude product (*R*)-*N*-((*S*)-2-aminopropyl)-2,4-dihydroxy-3,3-dimethylbutanamide (260 mg, 1.27 mmol, quantitative) as white solid.

**Rf:** 0.02 **(**5:1 EtOH in Heptane) **1H NMR** (500 MHz, DMSO-$D_6$) δ 8.01 (t, J = 5.7 Hz, 2H), 7.97 (s, 2H), 4.86 (s, 1H), 3.71 (s, 1H), 3.30 – 3.12 (m, 5H), 1.10 (d, J = 6.0 Hz, 3H), 0.79 (s, 3H), 0.78 (s, 3H).

3-(2-(3-(but-3-yn-1-yl)-3H-diazirin-3-yl)ethoxy)-*N*-((*S*)-1-((*R*)-2,4-dihydroxy-3,3-dimethylbutanamido)propan-2-yl)benzamide – Probe 2

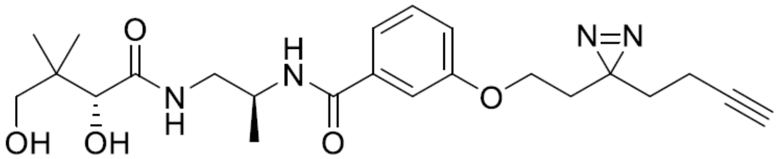

Under inert conditions (Ar) to a stirred solution was added 3-(2-(3-(but-3-yn-1-yl)-3H-diazirin-3-yl)ethoxy)benzoic acid (20 mg, 1 Eq, 77 µmol) **(Intermediate 6)** dissolved in dry DMF (1 mL) via syringe to a closed RBF followed by CDI (30 mg, 2.4 Eq, 0.19 mmol) dissolved in dry DMF (1 mL). The reaction mixture was analysed via MS. *(R)-N-*((*S*)-2-aminopropyl)-2,4-dihydroxy-3,3-dimethylbutanamide (30 mg, 1.9 Eq, 0.15 mmol) **(Intermediate 8)** was dissolved in dry DMF (1 mL) and triethylamine (71 mg, 97 µL, 9 Eq, 0.70 mmol) were added to the reaction mixture via syringe to a closed RBF and stirred overnight. The reaction was monitored via MS and TLC. The reaction mixture was concentrated in vacuo and purified using flash silica chromatography (0% → 10% MeOH in DCM) to afford pure product 3-(2-(3-(but-3-yn-1-yl)-3H-diazirin-3-yl)ethoxy)-*N*-((*S*)-1-((*R*)-2,4-dihydroxy-3,3-dimethylbutanamido)propan-2-yl)benzamide (4 mg, 9 µmol, 10 %) as a yellow oil.

**1H NMR** (500 MHz, CHLOROFORM-D) δ 7.38 – 7.26 (m, 4H), 7.06 (d, J = 7.1 Hz, 1H), 7.03 – 6.97 (m, 1H), 4.26 (dtp, J = 9.9, 6.6, 3.1 Hz, 1H), 4.02 (s, 1H), 3.86 (t, J = 6.2 Hz, 2H), 3.54 (ddd, J = 14.1, 9.7, 7.0 Hz, 1H), 3.39 (d, J = 7.6 Hz, 1H), 3.33 (ddd, J = 14.1, 5.5, 3.3 Hz, 1H), 2.06 (td, J = 7.5, 2.7 Hz, 2H), 1.99 (t, J = 2.7 Hz, 1H), 1.89 (t, J = 6.2 Hz, 2H), 1.72 (t, J = 7.5 Hz, 2H), 1.28 (d, J = 6.6 Hz, 3H), 0.91 (s, 3H), 0.86 (s, 3H). **13C NMR** (126 MHz, CHLOROFORM-D) δ 174.70, 167.42, 158.75, 135.53, 129.74, 119.40, 118.55, 112.86, 82.83, 78.50, 71.35, 69.36, 68.69, 62.82, 48.18, 47.81, 44.92 (d, J = 5.6 Hz), 39.07, 32.95, 32.72, 21.20, 20.90, 18.57, 13.38. **MS** (ESI) (m/z) calculated for $[M+H]^+$:445.24, found 445.15; calculated for $[M+Na]^+$: 467.23, found 467.15.

### NMR Spectra

#### 2,5-dioxopyrrolidin-1-yl 3-azido-5-ethynylbenzoate – Intermediate 2

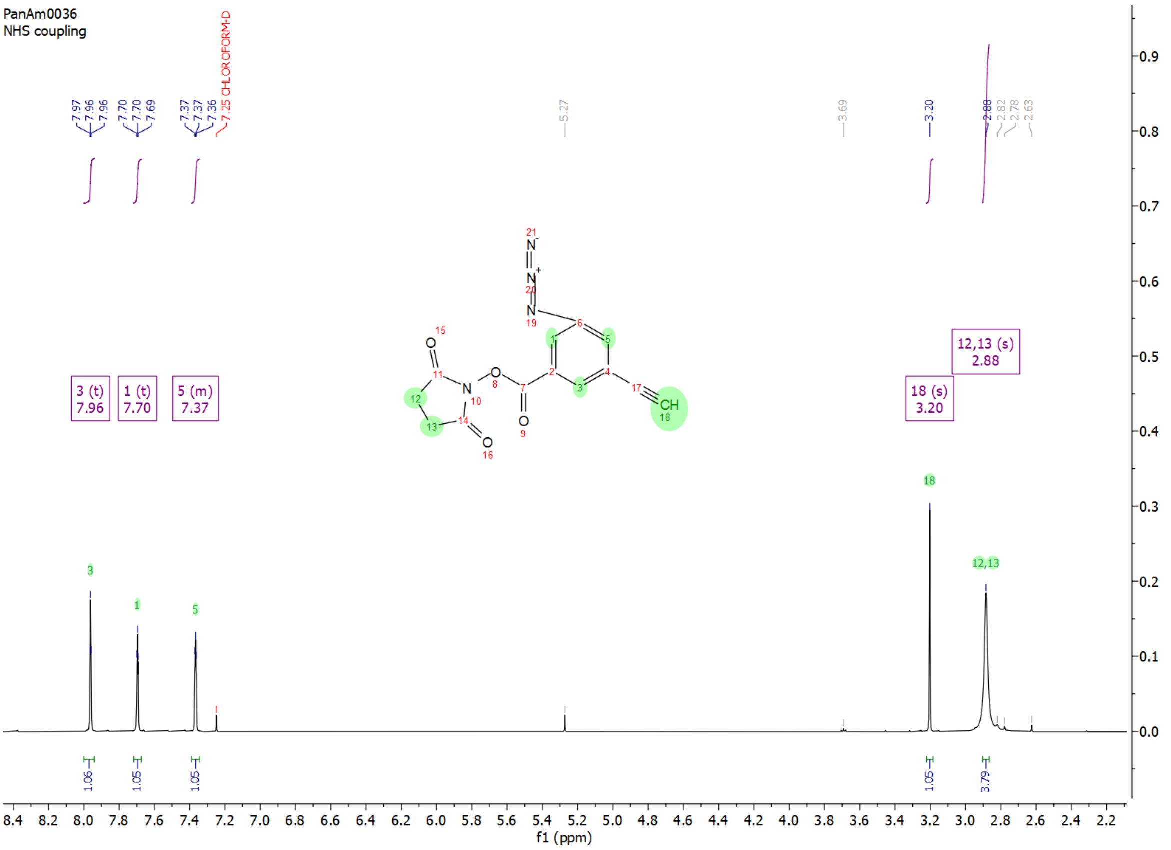


#### tert-butyl (S)-(2-(3-azido-5-ethynylbenzamido)propyl)carbamate – Intermediate 3

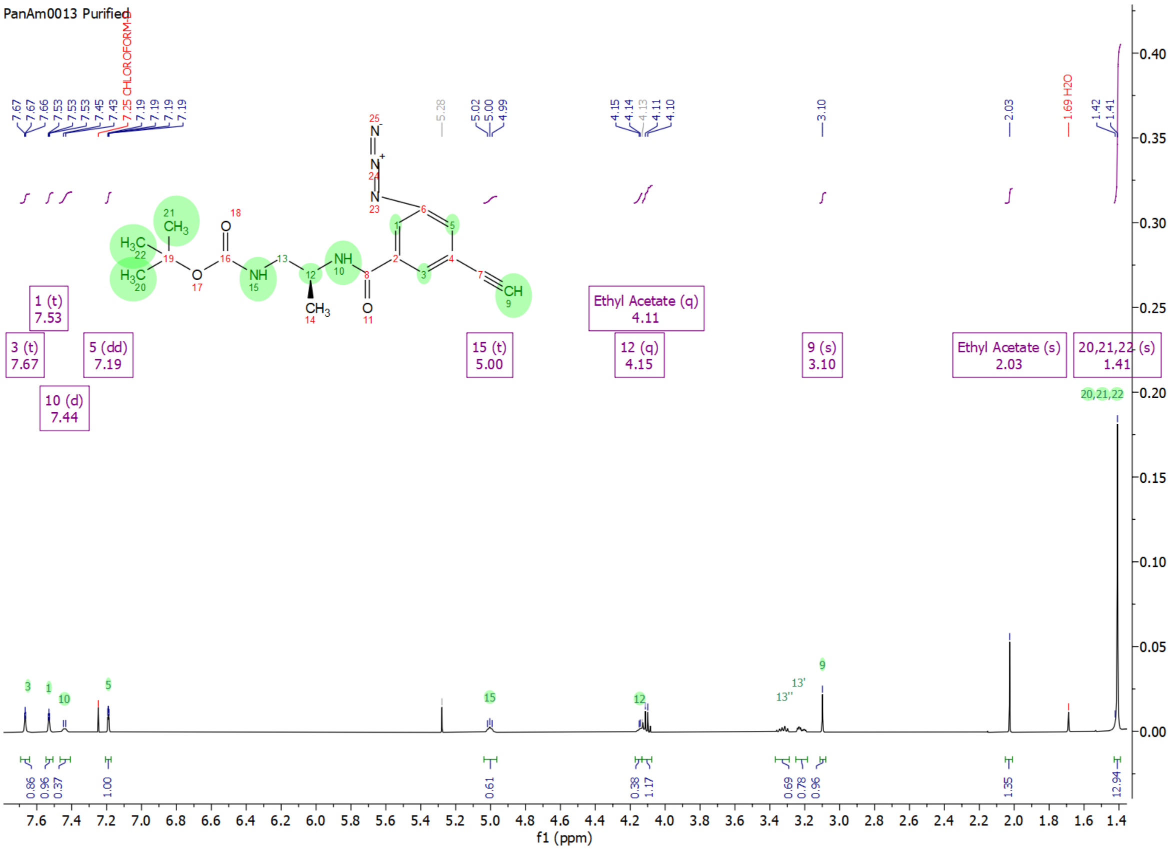

**(S)-N-(1-aminopropan-2-yl)-3-azido-5-ethynylbenzamide – Intermediate 4**

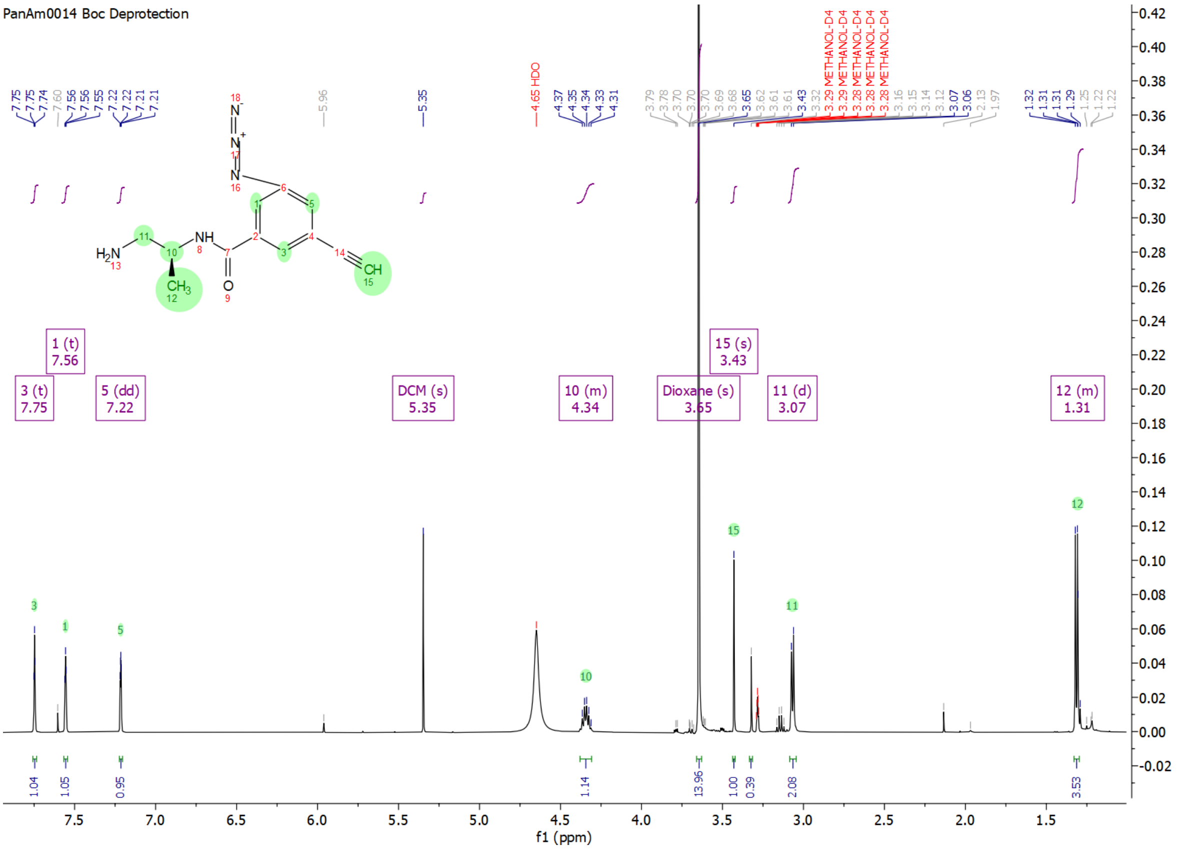

**3-azido-N-((S)-1-((R)-2,4-dihydroxy-3,3-dimethylbutanamido)propan-2-yl)-5- ethynylbenzamide – Probe 1**

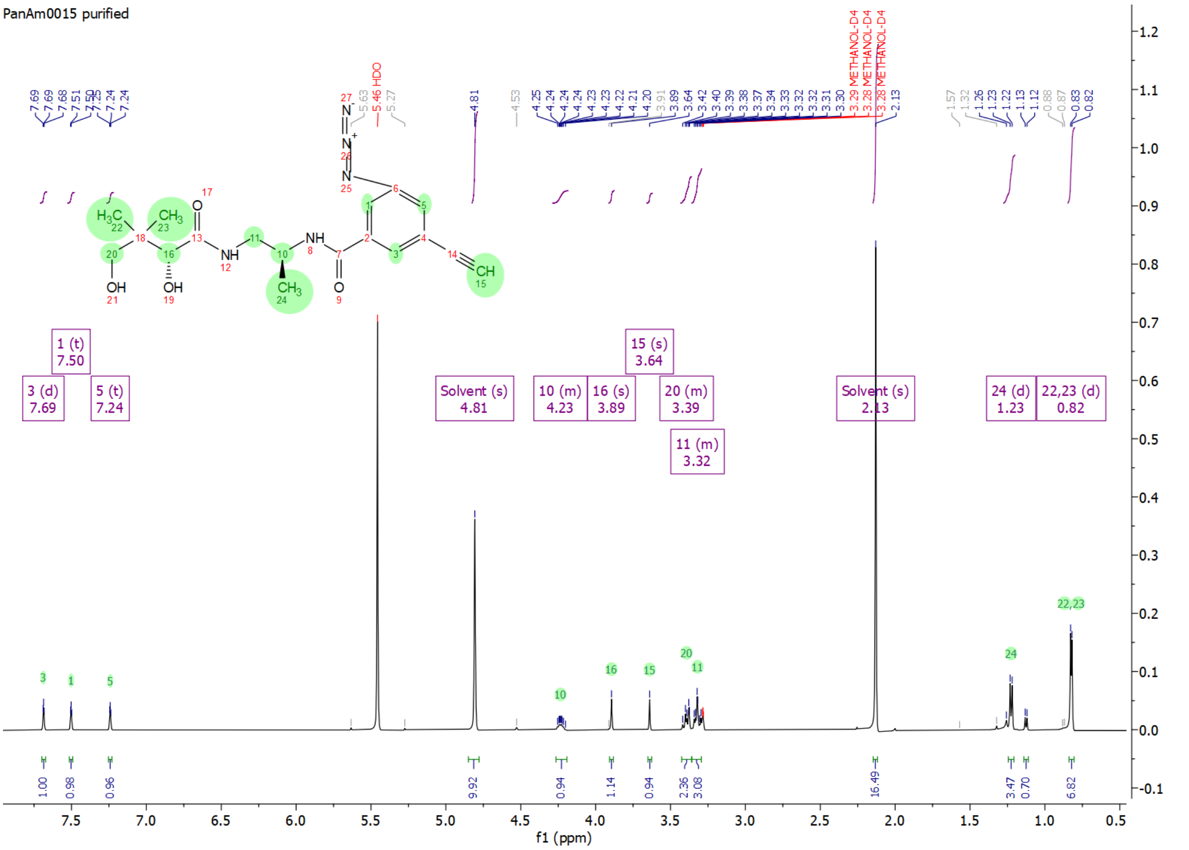


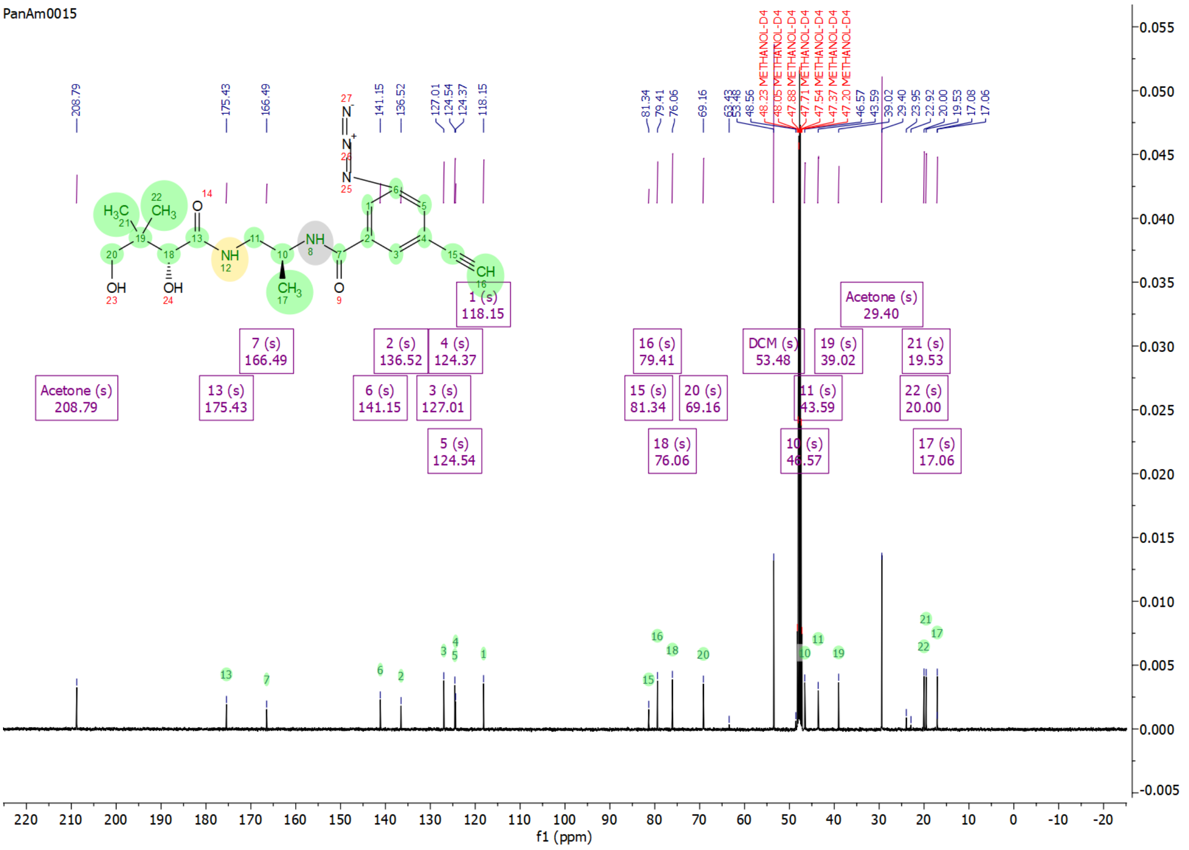

#### methyl 3-(2-(3-(but-3-yn-1-yl)-3H-diazirin-3-yl)ethoxy)benzoate – Intermediate 5

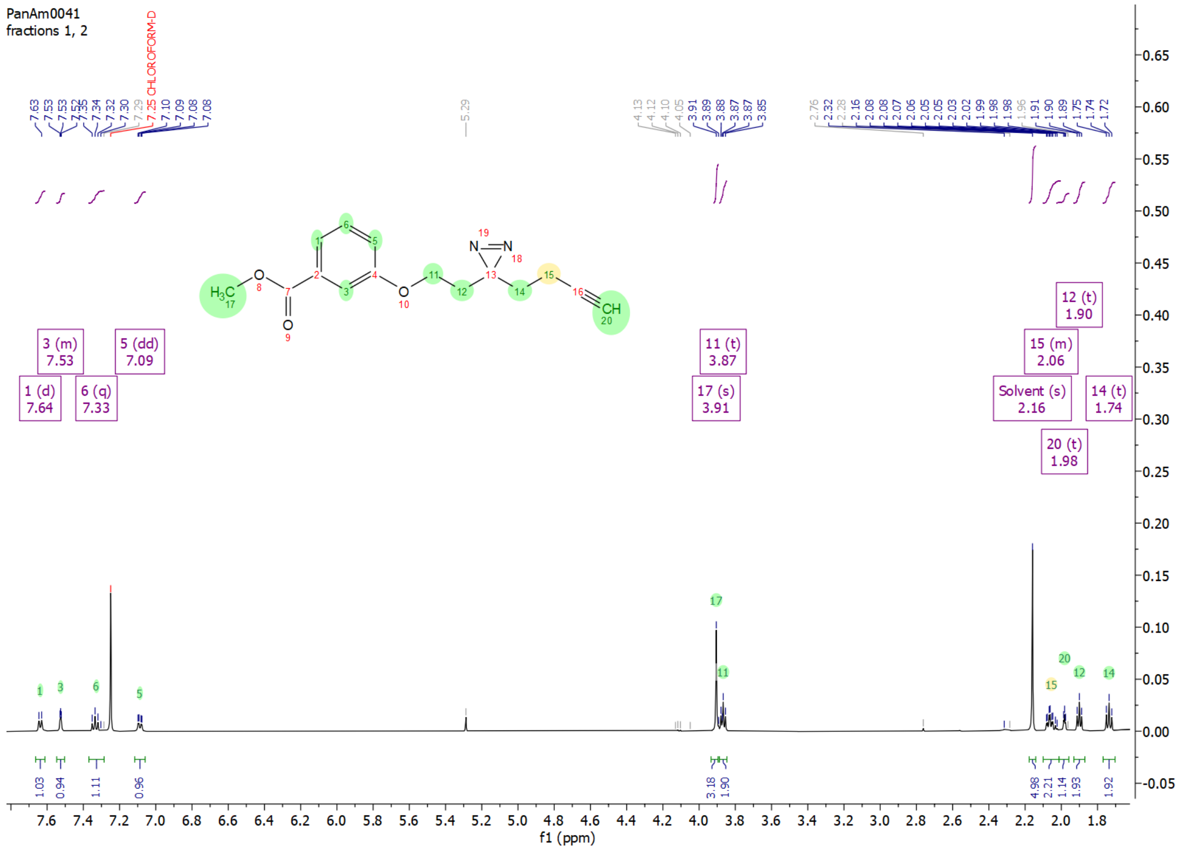


#### 3-(2-(3-(but-3-yn-1-yl)-3H-diazirin-3-yl)ethoxy)benzoic acid – Intermediate 6

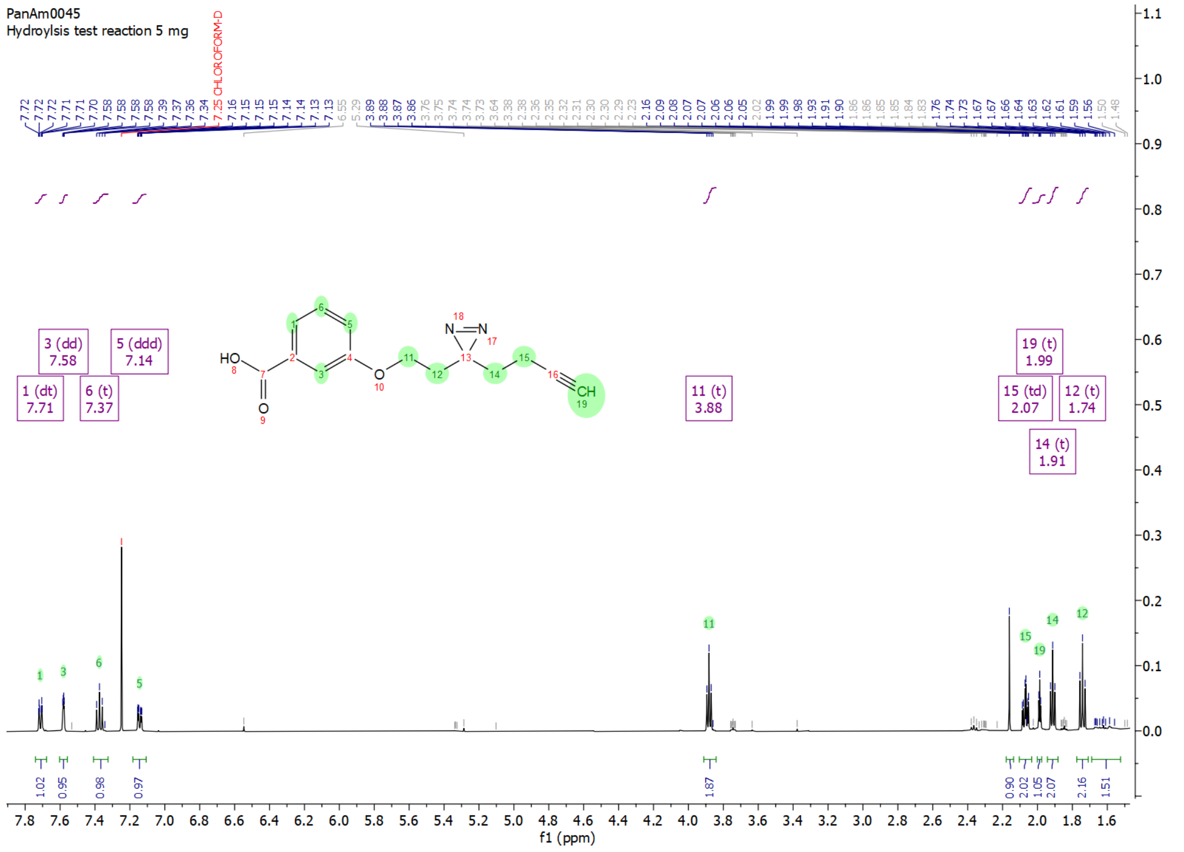

#### tert-butyl ((S)-1-((R)-2,4-dihydroxy-3,3-dimethylbutanamido)propan-2-yl)carbamate – Intermediate 7

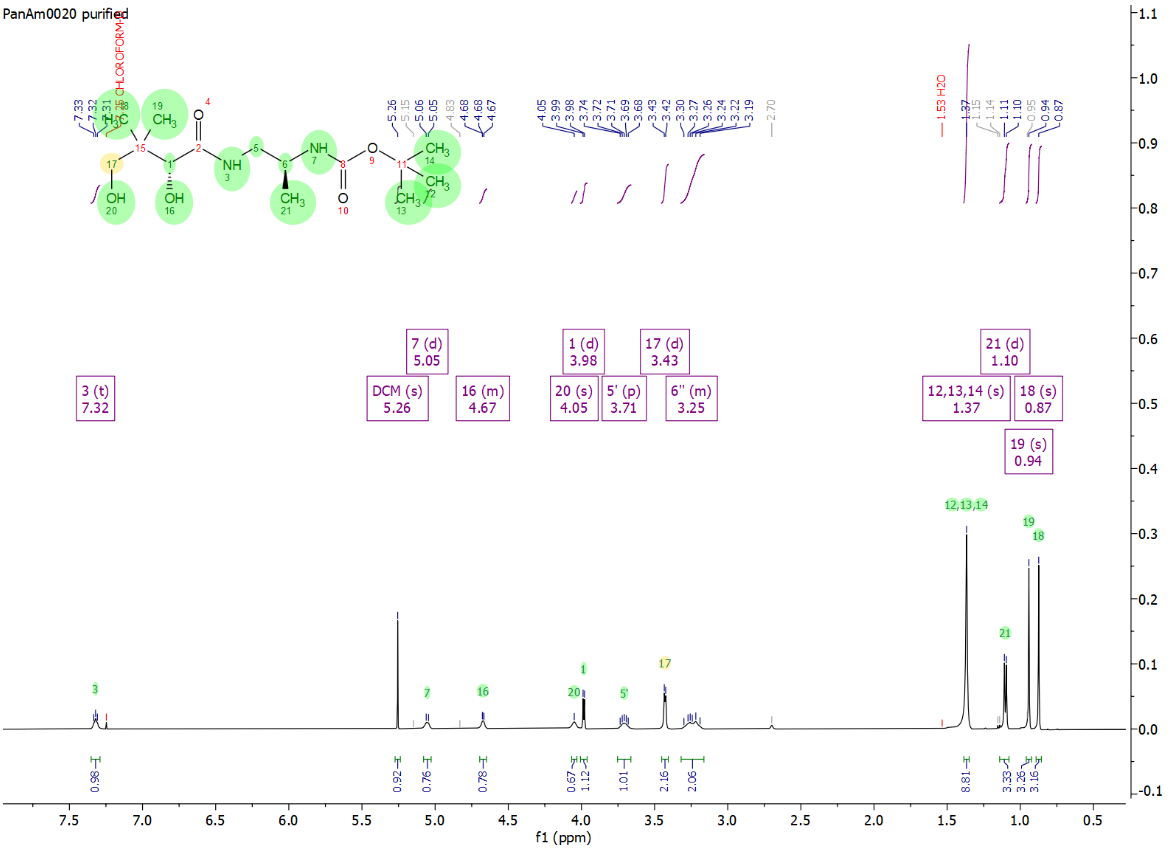


#### (R)-N-((S)-2-aminopropyl)-2,4-dihydroxy-3,3-dimethylbutanamide – Intermediate 8

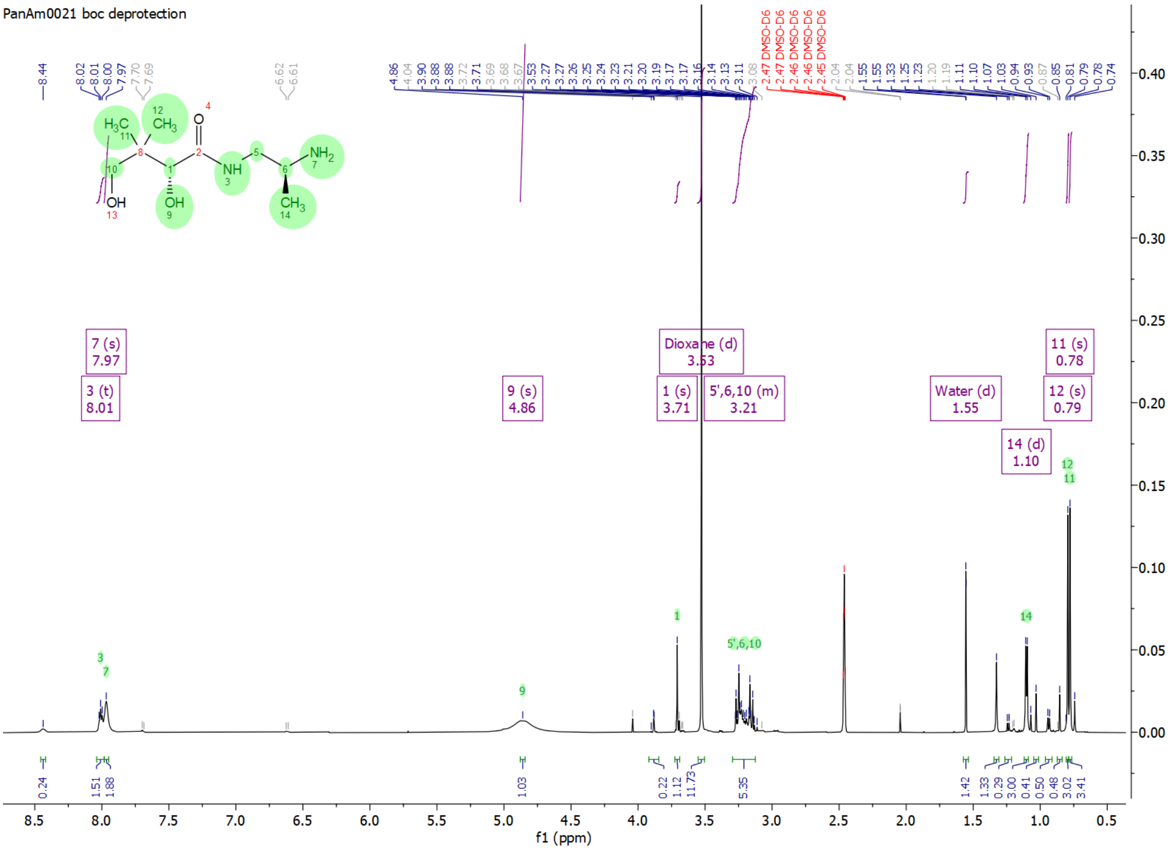

**3-(2-(3-(but-3-yn-1-yl)-3H-diazirin-3-yl)ethoxy)-N-((S)-1-((R)-2,4-dihydroxy-3,3-dimethylbutanamido)propan-2-yl)benzamide – Probe 2**

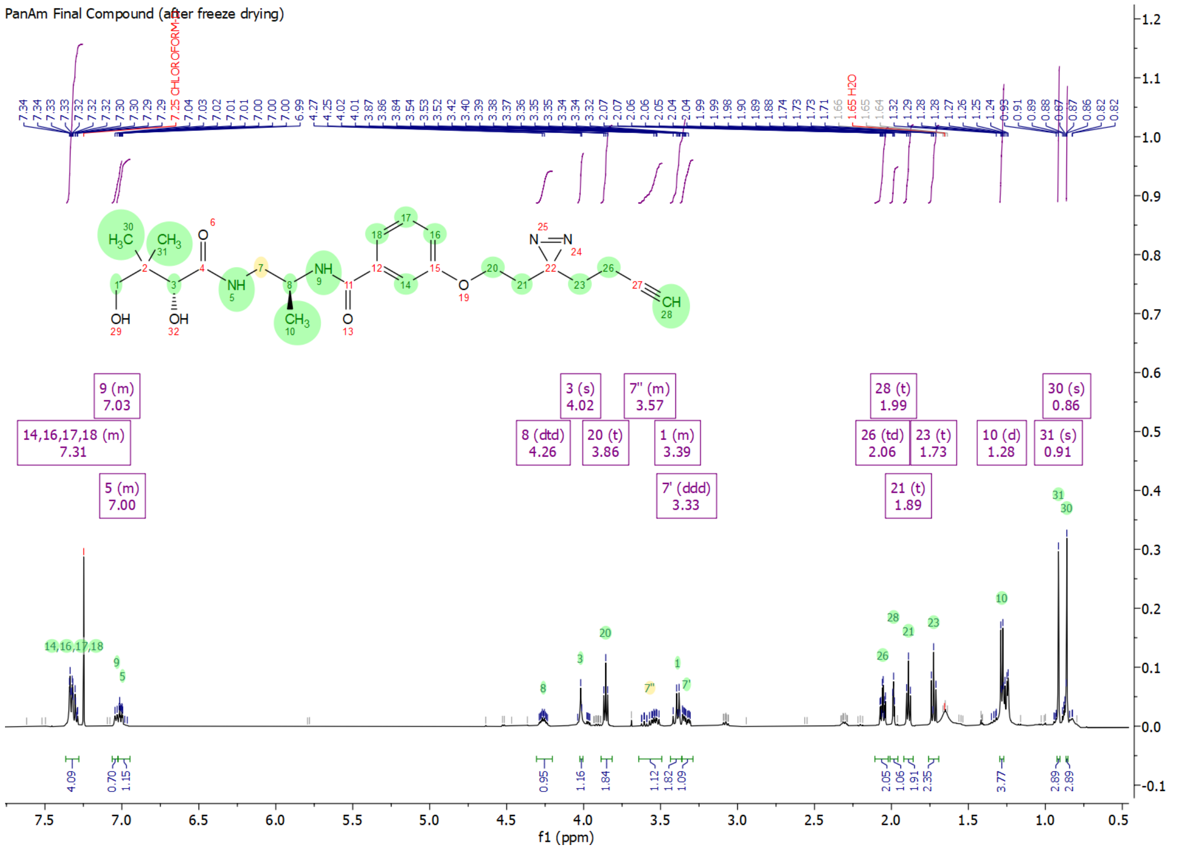


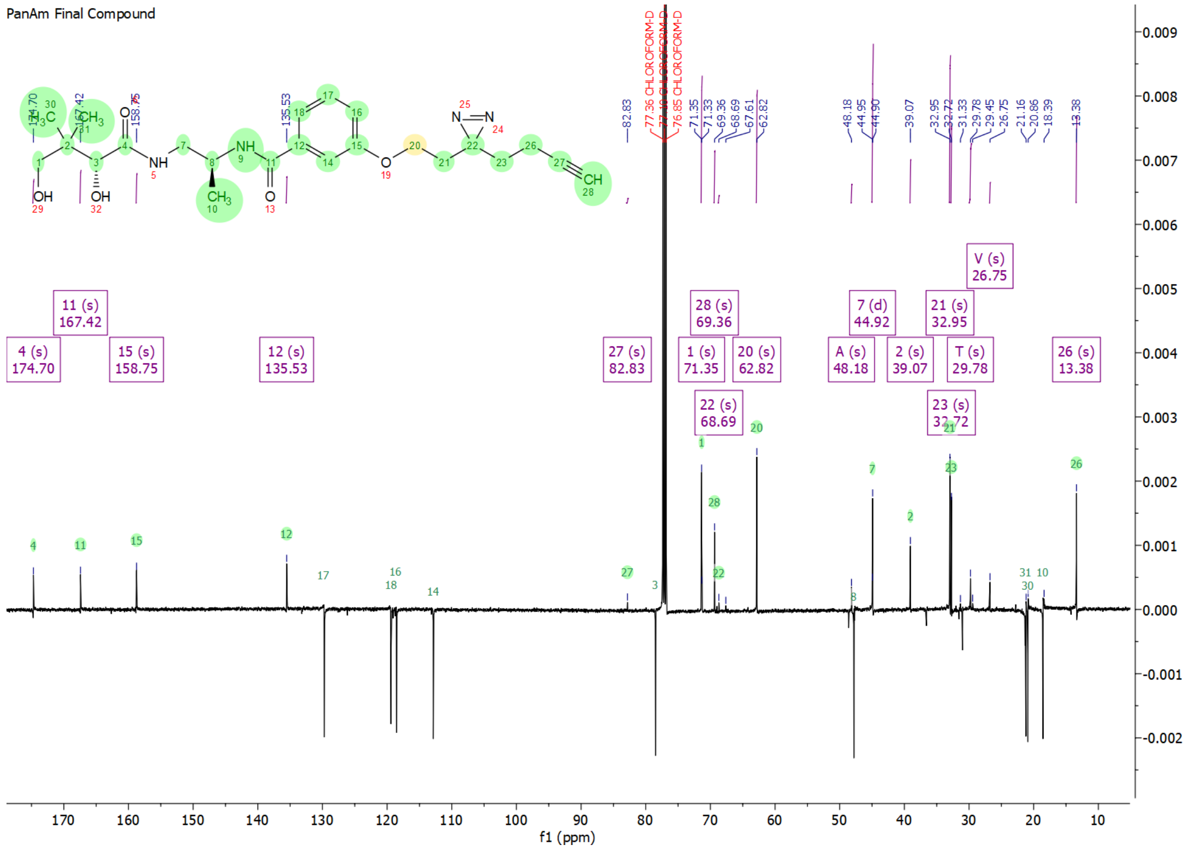

### Mass Spectrometry

**3-azido-N-((S)-1-((R)-2,4-dihydroxy-3,3-dimethylbutanamido)propan-2-yl)-5- ethynylbenzamide – Probe 1**

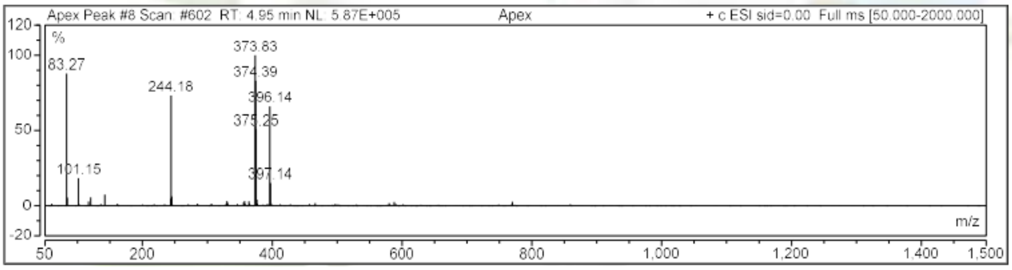


**3-(2-(3-(but-3-yn-1-yl)-3H-diazirin-3-yl)ethoxy)-N-((S)-1-((R)-2,4-dihydroxy-3,3-dimethylbutanamido)propan-2-yl)benzamide – Probe 2**

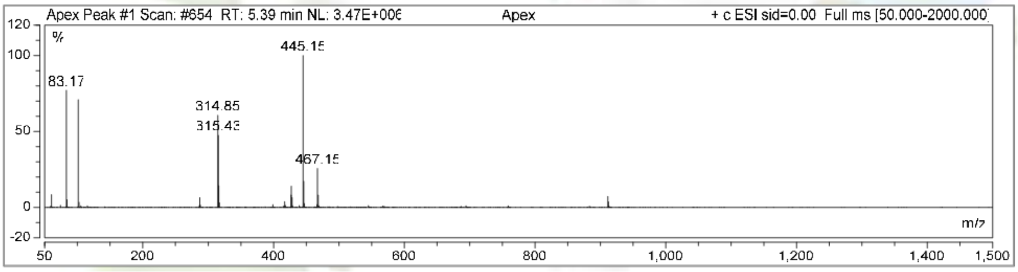
